# Effect of human behavior on the evolution of viral strains during an epidemic

**DOI:** 10.1101/2021.09.09.459585

**Authors:** Asma Azizi, Natalia L. Komarova, Dominik Wodarz

**Affiliations:** Department of Mathematics, Kennesaw State University, Marietta, GA 30060; Department of Mathematics, University of California Irvine, Irvine CA 92697; Department of Population Health and Disease Prevention Program in Public Health, Susan and Henry Samueli College of Health Sciences, University of California, Irvine CA 92697

**Keywords:** Mandated social distancing, Self-regulated social distancing, Network, Viral evolution, Symptomatic variant, Asymptomatic variant

## Abstract

It is well known in the literature that human behavior can change as a reaction to disease observed in others, and that such behavioral changes can be an important factor in the spread of an epidemic. It has been noted that human behavioral traits in disease avoidance are under selection in the presence of infectious diseases. Here we explore a complimentary trend: the pathogen itself might experience a force of selection to become less “visible”, or less “symptomatic”, in the presence of such human behavioral trends. Using a stochastic SIR agent-based model, we investigated the co-evolution of two viral strains with cross-immunity, where the resident strain is symptomatic while the mutant strain is asymptomatic. We assumed that individuals exercised self-regulated social distancing (SD) behavior if one of their neighbors was infected with a symptomatic strain. We observed that the proportion of asymptomatic carriers increased over time with a stronger effect corresponding to higher levels of self-regulated SD. Adding mandated SD made the effect more significant, while the existence of a time-delay between the onset of infection and the change of behavior reduced the advantage of the asymptomatic strain. These results were consistent under random geometric networks, scale-free networks, and a synthetic network that represented the social behavior of the residents of New Orleans.

## 1 Introduction

Epidemic spread of infectious diseases is a topic that has received much attention among computational modelers, see e.g. [1, 2, 3, 4, 5]. One important aspect of this process is the rise and spread of mutant variants of the pathogen [6, 7, 8, 9, 10, 11]. For example, in a spatially expanding epidemic, it was shown that less virulent strains will dominate the periphery while more virulent strains will prevail at the core [12]. It has also been observed that in epidemic models where infection events happen on an interaction network, evolutionary dynamics of the pathogen change depending on the structure of the network [13, 14, 15, 16]. It has been shown, for example, that heterogeneities in contact structure (i.e. network degree) may accelerate the spread of a single disease, and at the same time slow down the rise of a rare advantageous mutation under susceptible-infected-susceptible (SIS) infection dynamics [17]. In the context of spatial networks with host migration, it was reported that the spatial network structure may have important effects on the transient evolutionary dynamics during an epidemic [18]; in particular, the front and the rear of the expanding epidemic are expected to be phenotypically different. Pinotti et. al. [19] studied the influence of the social network structure on competition dynamics of strains (with identical parameters) that are spread via a stochastic SIS model on the network. It was found that network structure can affect the ecology of pathogens: in a more heterogeneous network, a reduction in the number of strains and an increase in the dominance of one strain were observed, while strong community structure in the social network increased the strain diversity.

Another relevant characteristic of epidemic dynamics that has been investigated is the effect of human behavior on disease spread, see e.g. [20, 21, 22, 23]. Different aspects of human behavior have been considered, including relational exchange (e.g. replacement of sick individuals by healthy ones in the workplace) [24], people’s hygiene [25], voluntary vaccination and vaccination compliance [26], “risky” versus “careful” individual behavior [27, 28, 29, 30, 31], and the related concept of social distancing. Social distancing is a change of behavior that can roughly be classified into (1) self-regulated (or spontaneous) where individuals may choose to limit their contacts based on information that they receive or on their personal beliefs [32, 33, 31, 34, 35, 36]; and (2) mandated (public), where the decrease in social contacts is regulated centrally and affects either the entire population or certain subpopulations [37, 38]. The COVID19 pandemic has triggered much research into the role of social distancing in viral spread, especially because before the advance of vaccination, non-pharmaceutical intervention (NPI) measures were the only way of intervention available [39]. NPI policies have taken a variety of forms such as extreme lock-downs, school closure, road and transit systems restrictions, and mandatory isolation/ quarantine [40], see e.g. [41, 42, 43, 44, 45, 46, 47, 48, 49, 50, 51] on the effects of mandated social distancing on SARS-CoV-2 spread. In a recent paper [52] the authors considered the combination of both mandated and self-regulated types of social distancing, and studied their effect on the outbreak threshold of an (asymptomatic) infectious disease.

In this paper we explore the role of mandated and self-regulated social distancing on viral evolution. The focus of this study is the co-evolution of two types of a pathogen, the resident, more symptomatic, pathogen, and an emerging, less symptomatic (or asymptomatic), variant. The two may or may not differ in their infectivity properties, but because they present differently, they will trigger different behavior of the individuals, which may result in different levels of self-regulated social distancing. As a result, the less symptomatic variant may experience a selective advantage. We will use the usual framework of the susceptible-infectious-removed (SIR) model on networks, and investigate how the network structure (including random networks of different types and a synthetic network representing social interactions of real individuals) modifies the co-dynamics of the two viral strains.

## 2 Methods

The model includes the infection dynamics transmission and intervention strategies. It is assumed that the disease spreads within a Susceptible–Infected–Removed (SIR) framework. Dynamics take place on a network, and three different network types are studied.

### 2.1 Network structure

We assume that the epidemic spreads on a network of size *N*, where each node represents a person, and the edges represent interactions. Here we study two types of random, unweighted networks: the random geometric network, and the scale-free network (with *N* = 10, 000 nodes). Each of these networks represents a different type of abstraction that retains certain features of human interactions. In addition to these two types of random networks, we also studied disease spread on a real-world synthetic network of a much larger size (*N* = 150, 000), where the edges are weighted by the time the two individuals spend together. This synthetic network was constructed based on interaction data of people in New Orleans [53, 54].

#### Random Spatial-Geometric Network

This network is constructed by placing *N* points in a unit square and connecting only the points that are within a prescribed Euclidean distance, *r*, from each other. Such networks are characterized by a strong local structure and clustering properties, and have been studied extensively in the literature [55, 56]. Such networks could represent local social contacts of individuals in the absence of any long-range connections.

#### Scale-free Network

This network is characterized by a power law degree distribution. As a result, while most individuals only have a limited number of contacts, there are “super-spreaders” of very high degrees [57, 58]. Examples of applications of such networks are the number of sexual partners in a college environment [59] or the network of a city with buildings (nodes) and flows of people as connecting edges [60].

We use Networkx open software platform [61] to generate Spatial-Geometric random networks in dimension 2 and and distance threshold *r* = 0.02. We also use the Barabási–Albert preferential attachment model in Networkx to generate scale-free networks with degree distribution *P*(*k*) ∼ *k*^−2.11^. The random networks have the same size and average degree, but they differ in terms of their degree distributions and other properties, since they have different structures.

Each of these networks has advantages and disadvantages when used to model epidemic spread in populations. Random spatial-geometric networks successfully model clustering properties of human interactions but do not include long-range connections or superspreaders. Superspreaders are a natural part of scale-free networks, but the latter network type has no clustering or neighborhood structure. For these reasons we perform all the analyses for different network types, to investigate whether observed phenomena depend on any particular network properties. Finally, we implement the most realistic network in the study, the New Orleans synthetic network, which is described below.

#### Real World Network

Our real world network is based on the synthetic data generated by Simfrastructure [53, 54] for *N* = 150, 000 synthetic people residing in New Orleans. Simfrastructure is a high-performance, service-oriented, agent-based modeling and simulation system for representing and analyzing interdependent infrastructures. In the New Orleans network, each edge *ij* between two nodes *i* and *j* is weighted by *ω*_*ji*_, which represents the strength of connectivity between *i* and *j*, and reflects the type of connection as well as the amount of time the two individuals spend with each other.

### 2.2 SIR model on a network for two virus strains

In our stochastic Susceptible–Infectious-Removed (SIR) model superimposed on the network, an individual *i* at time *t* is either susceptible to being infected, infected, or removed from the infection because of recovery or death. During a time-interval Δ*t*, an infected individual can infect any of their susceptible neighbors (that is, susceptible individuals connected with them by an edge). We denote by *β* the infection rate per edge, such that during time Δ*t*, the probability that a susceptible individual *j* will be infected by an infected neighbor *i* is given by *βω*_*ji*_Δ*t*. (Note that for the random spatial and scale-free networks, we will use *ω*_*ji*_ = 1). For each infected individual, a recovery event occurs during the time-interval Δ*t* with a probability *γ*Δ*t*, or a death event occurs with a probability *δ*Δ*t*, and we refer to the rate of death or recovery as the rate of removal, *ρ* = *γ* + *δ*.

We assume the existence of two distinct variants (strains) of the virus, which we denote by *V*_1_ and *V*_2_. Our model incorporates permanent cross-immunity for either viruses, that is, if an individual is infected by virus *k*, then they are immune to virus *k*′ for *k*′ ≠ *k* during their infection and after recovery (here *k, k*′ ∈ {1, 2}). We further assume that an individual infected with virus *k* can only induce infection with virus *k*, that is, we do not consider spontaneous mutations from one type of virus to the other.

Unless noted otherwise, the two virus strains are assumed to have identical parameters, that is, the same values of *β, δ*, and *γ*. The only difference between the two strains is that one (*V*_1_) causes a symptomatic disease, while the other (*V*_2_) is asymptomatic. This gives rise to differences in people’s behavior, as described in the next subsection. Later on, we consider scenarios in which symptomatic infection is coupled to a higher viral infectivity.

For initialization, we start the epidemic by randomly infecting one individual with *V*_1_. We then advance the simulation until the epidemic grows to 0.1% *V*_1_-infected individuals. At this time we introduce the next randomly generated newly infected case as a *V*_2_ infection; this represents a single mutation event of the resident strain.

At this point, we reset the time to zero and use this state as the initial condition to study the virus co-dynamics in the absence of any further mutant generation.

Simulation speed depends on the size of time-step Δ*t*, so it is desirable to pick the largest value for Δ*t* such that the simulations exhibit reasonable convergence accuracy, see also [62]. We have implemented the program for the null scenario (no social distancing) with Δ*t* values representing 1 day, 1 hour, and 1 minute, and while results differed significantly between Δ*t* = 1 day and Δ*t* = 1 hour, the the result for Δ*t* = 1 hour and Δ*t* = 1 minute were almost identical. Therefore, we chose Δ*t* = 1 hour for our simulations in this study.

### 2.3 Social distancing strategies

We model two types of social distancing (SD) strategies: (1) mandated SD implemented by the government, and (2) self-regulated SD.

Mandated SD is implemented as follows: when the prevalence of virus (i.e. the fraction of infected individuals among the population) reaches a fixed threshold *ψ*, all individuals start practicing temporary social distancing. To this end, the fraction *σ*_*M*_ of all the edges in the network are removed for *τ*_*M*_ consecutive days; connections to be removed are chosen randomly.

Self-regulated SD is also implemented only if the number of infections has reached the threshold prevalence *ψ*. If an individual has at least one neighbor that is symptomatically infected with *V*_1_ (after a delay *τ*_*s*_ following infection), they remove fraction *σ*_*S*_ of their connections. The connections to be removed are chosen randomly, and remain cut for as long as there is a symptomatically infected neighbor.

It is possible that fraction *σ*_*S*_ or *σ*_*M*_ of connections is a non-integer number, *K*. In this case, if [*K*] stands for *K*’s integer part, [*K*] + 1 connections are removed with probability *K* − [*K*], and [*K*] connections are removed otherwise.

### 2.4 Parameter values

The definitions of all the variables and parameters of the proposed model are given in the table 1. The parameter values have been chosen to be realistic for respiratory infections and are specified in the figure legends. Under these parameters, the basic reproduction number comes out to be between 2 and 3 for the examples considered.

**Table 1:** Parameter and state variable definitions and notations.

|  | Notation | Description | Unit |
| --- | --- | --- | --- |
| Network<br>Parameters | $N$ | Number of nodes in the network | People |
|  |  | Spatial network | – |
|  |  | Scale-free random network | – |
|  |  | Real world network | – |
| | $\omega_{ij}$ | The connectivity level between two neighbors $i$ and $j$ | 1 |
| | $\bar{C}$ | Average number of contact per time for random networks | Contact/time |
| Infection<br>Parameters | $\beta_k$ | Prob. of $V_k$ transmission per contact per time | 1/contact |
| | $\rho$ | Per time removal (death or recovery) probability from virus $k$ | 1/time |
| | $\tau_s$ | Time-period between getting infection and showing the symptoms for $V_1$ infected cases | Time |
| Intervention<br>Parameters | $\psi$ | Prevalence threshold: Infection prevalence to start SD | 1 |
| | $\sigma_M$ | Mandated SD: fraction of removed contacts | 1 |
| | $\sigma_S$ | Self-regulated SD against $V_1$: fraction of removed contacts | 1 |
| | $\tau_M$ | Duration of mandated SD | Time |

To estimate the reproduction number ℛ_0_, starting with randomly selected individual as initial infected case, we count the number of neighbors who get infected from them during their infection period. We repeat this process for a large number of independent simulations, seeding different initial infected individuals. Intervention parameters will change based on different scenarios explored here, and are specified in figure legends.

## 3 Results: positive selection of the asymptomatic strain on different networks

Here we explore the consequences of behavioral changes (self-regulated social distancing) on the spread of an asymptomatic viral strain. First this is done by using two types of abstract random networks, the scale-free and the random spatial network. Both types of random networks have some features resembling different aspects of human social networks. Then we show how similar scenarios play out on a more realistic network that emulates the behavior of a real-life population of New Orleans.

### 3.1 Self-regulated social distancing selects for an asymptomatic strain

In our model, individuals in the population exercise self-regulated SD if members of their circle become symptomatically infected (that is, become infected with *V*_1_). To explore the consequence of this behavior on the evolutionary dynamics of asymptomatic virus variants (*V*_2_), we ran simulations where such a mutant was introduced as a minority in the initial stages of the epidemic, see figure 1. We explored the dynamics on two different networks: scale-free (left panels) and spatial (right panels); the trajectories presented are averages over 5000 independent simulations. We present four different degrees of self-regulated SD: *σ*_*S*_ = 0 (a control case where *V*_2_ is indistinguishable from *V*_1_ in the model, and no selection is expected), *σ*_*S*_ = 0.2 (low-degree self-regulated SD), *σ*_*S*_ = 0.4 (moderate self-regulated SD), and *σ*_*S*_ = 0.7 (high-degree self-regulated SD). As time goes by and the epidemic spreads, we plot the prevalence of each virus (panels (1a) and (1b)), and also follow the relative share of *V*_2_, that is 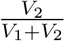(panels (1c) and (1d)).

**Figure 1:**
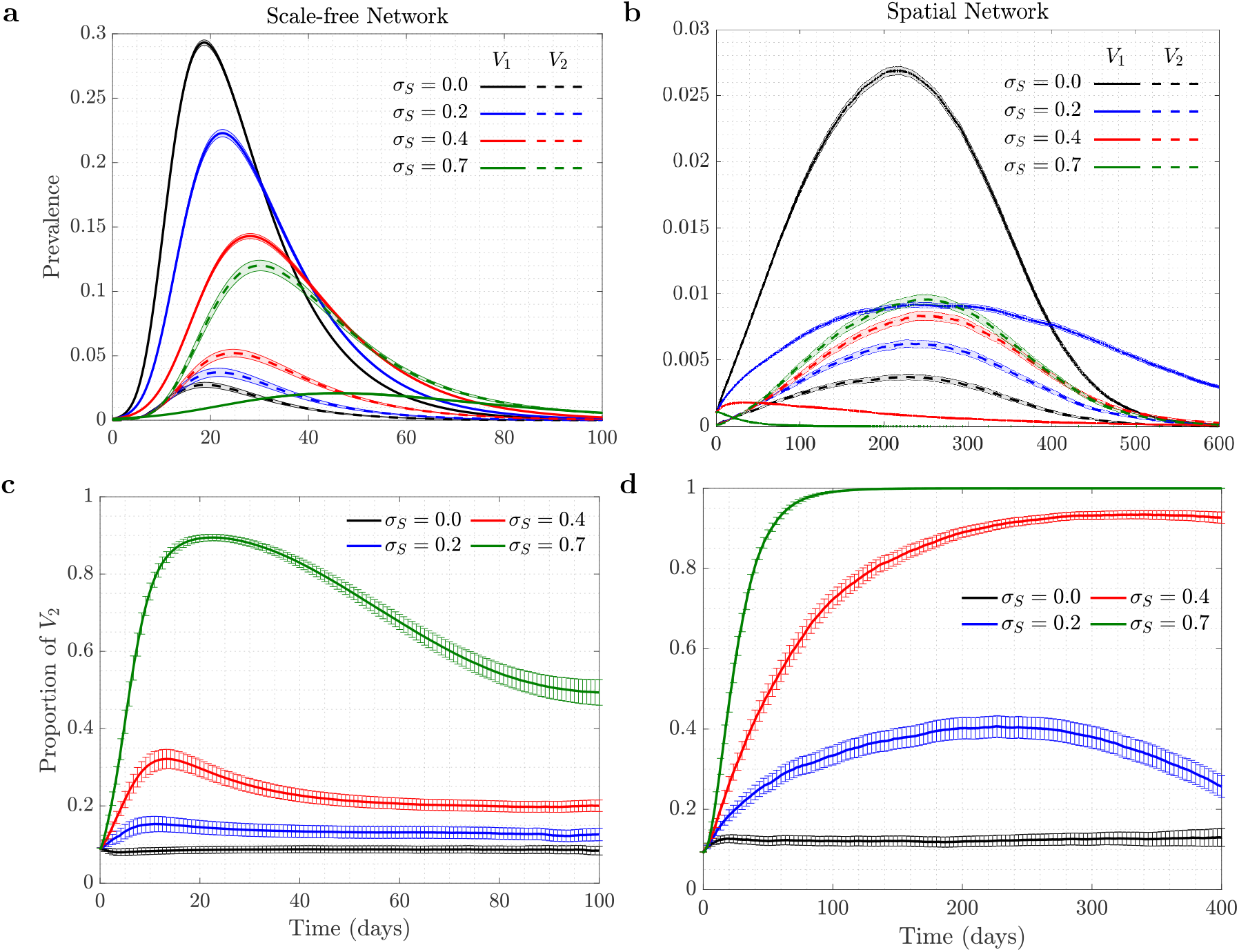
The role of self-regulated SD in the spread of viruses. Time series are shown for four scenarios of no (*σ*_*S*_ = 0, black), low (*σ*_*S*_ = 0.2, blue), moderate (*σ*_*S*_ = 0.4, red), and high (*σ*_*S*_ = 0.7, green) self-regulated SD, in the absence of mandated SD. Scale-free (left) and spatial (right) networks of 10, 000 individuals and average degree 10 are used. Panels (a, b) plot are the prevalence of *V*_1_ (solid) and *V*_2_ (dashed); panels (c,d) show the proportion of *V*_2_ (*V*_2_/(*V*_1_ + *V*_2_)). The rest of the parameters are *γ* + *δ* = 0.1 per day, *ψ* = 0.0012, *β*_1_ = *β*_2_ = 0.028 per day per contact for scale-free and *β*_1_ = *β*_2_ = 0.037 per day per contact for spatial network (corresponding to ℛ_0_ = 2.5). Means and standard errors are shown for 5000 stochastic realizations.

In the absence of self-regulated SD (black lines in panels (1a) and (1b)), the epidemic on the two networks looks different despite similar ℛ_0_ parameters: infection burns through the scale-free network faster and reaches a higher infection peak, while in the case of the spatial network it lasts longer at relatively low levels.

Under zero self-regulated SD (black lines in panels (1c) and (1d)), as expected, the proportion of *V*_2_ remains approximately constant throughout the course of the epidemic, although we do observe an initial increase in the abundance of *V*_2_ in the spatial network. This initial increase is due to a somewhat “advantageous” initial location of the *V*_2_ infection. In the spatial network, it gets placed on the “outskirts” of the growing infected neighborhood, which results in a larger mean number of uninfected neighbors that *V*_2_-infected individuals have compared to *V*_1_-infected individuals. This initial increase of the proportion of *V*_2_ is therefore due to the initial placement and does not represent an ongoing selection.

A different pattern is observed in the presence of self-regulated SD: the proportion of *V*_2_ infected individuals increases well beyond the initial boost. This effect is stronger for a larger extent of self-regulated SD (compare green (*σ*_*S*_ = 0.7) to red (*σ*_*S*_ = 0.4) to blue (*σ*_*S*_ = 0.2) lines in the bottom panels of figure 1). The exact extent to which the fraction of *V*_2_ increases in the course of the epidemic depends, besides *σ*_*S*_, on the network size and type. Larger networks will result in a larger increase in *V*_2_ fraction, simply because they experience a larger and longer epidemic, and *V*_2_ will have a longer time to gain on *V*_1_ before the epidemic runs out of targets (not shown); a similar result can be demonstrated by using an ODE model of an SIR infection with two viral strains, see supplementary material.

We note a significant difference in the amount of gain experienced by the asymptomatic strain under scale-free (panel (c)) and spatial (panel (d)) networks. Self-regulated SD results in much more effective protection on a spatial network, because if an individual has an infected neighbor, they are likely to have more than one infected neighbor, and self-regulated SD induced by one of the neighbors will work against future infections in the vicinity. This results in a much larger force of selection experienced by the asymptomatic strain on a spatial network, compared to the case of scale-free network, which does not have a community structure. More details are presented in supplementary material.

### 3.2 Advantage mediated by self-regulated SD can off-set a fitness cost of the asymptomatic strain

Figure 2 explores a scenario where the asymptomatic mutant, *V*_2_, has a fitness cost compared to the resident virus, *V*_1_, which is manifested through a reduction in the probability of transmission parameter. We can see that although having a small disadvantage in *β*_2_ reduces the fraction of *V*_2_, we still observe a rise in the prevalence of *V*_2_ caused by self-regulated SD against symptomatic cases. In other words, the behavior-based selection mechanism can work even in the presence of a degree of disadvantage in the transmissibility of the mutant compared to the resident type. We observe that even in the presence of a significant disadvantage of virus *V*_2_, self-regulated SD can provide enough pressure to lead to positive selection of the asymptomatic virus. Again, we note a difference in the force of selection for the asymptomatic strain under scale-free and spatial networks. In the case of a scale-free network, (figure 2(a)) a 15% disadvantage of *V*_2_ almost completely eliminates any advantage gained through self-regulated SD. In the case of a spatial network (figure 2(b)), an asymptomatic strain with a 15% fitness costs still rises to almost 90% in the population.

**Figure 2:**
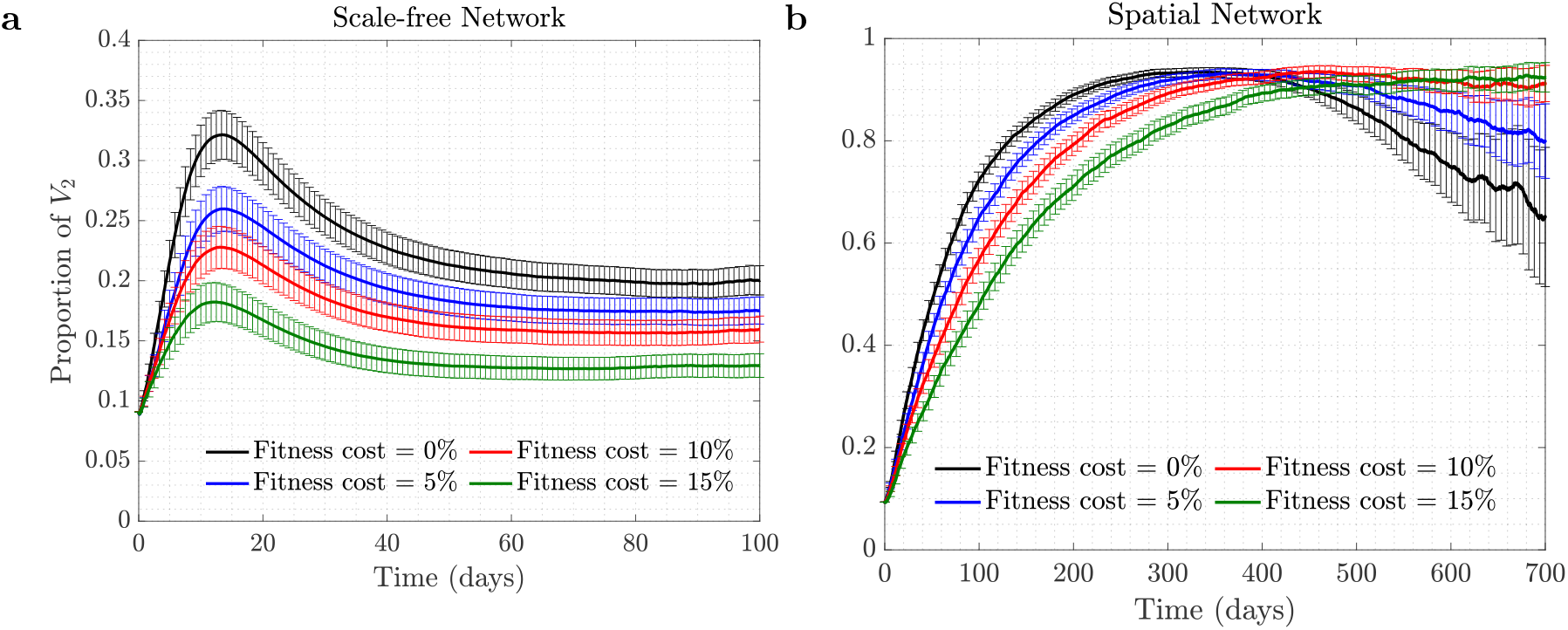
Selection for *V*_2_ in the presence of a fitness cost. Time series of proportion of *V*_2_ under moderate self-regulated SD, *σ*_*S*_ = 0.4 (and with *σ*_*M*_ = 0), are shown for 0% fitness cost (*β*_2_ = *β*_1_, black), 5% fitness cost (*β*_2_ = 0.95*β*_1_, blue), 10% fitness cost (*β*_2_ = 0.9*β*_1_, red), and 15% fitness cost (*β*_2_ = 0.85*β*_1_, green), for (a) scale-free and (b) spatial networks. All the other parameters are as in Figure 1.

### 3.3 Mandated social distancing makes selection stronger

Next, we explored the consequence of mandated SD implementation on the selection for the asymptomatic strain. Mandated SD affects transmission of both viral strains equally, and it is not immediately clear whether the presence of mandated SD can modify the dynamics and change the advantage experienced by *V*_2_ through self-regulated SD. Figure 3 assumes the presence of self-regulated SD at an intermediate level, and shows that increasing the level of mandated SD increases the positive selection pressure experienced by the asymptomatic strain. As a function of time, the fraction of *V*_2_ grows at the same rate for all levels of mandated SD (that is, the initial slope of the fraction is defined by the level of self-regulated SD and independent of the mandated SD). The dynamics are however different at later times, where the peak of the *V*_2_ fraction is higher (and is reached later) for higher levels of mandated SD. The reason for this event is that increasing mandated SD results in a reduction in the reproduction number, ℛ_0_, which generally leads to a longer, lower-level epidemic, so the fitter virus (*V*_2_) has a longer time to expand relative to its symptomatic counterpart. Once the epidemic is on the decline, the fraction of *V*_2_ decreases (see supplementary material; the same trend is observed for the spatial network on a longer time-scale, not shown). Figure 3 shows that the fraction of the asymptomatic strain among all infected individuals increases with the level of mandated SD. A similar result is demonstrated in an ODE SIR model for two viral strains, see supplementary material, figure 1(a). In the ODE model, we could not directly include a network structure or details of mandated or self-regulated SD. Instead, to gain indirect insights into the system of interest, we investigated the co-dynamic of two strains in a population with complete mixing, where strain *V*_2_ was characterized by a larger fitness compared to strain *V*_1_. This was achieved explicitly by increasing *V*_2_’s infectivity, and represents fitness differences due to self-regulated SD. Keeping the relative fitness of the strains fixed, we reduced the overall fitness of both strains (this mimics degrees of mandated SD, which reduces the infectivity of both strains equally). It was shown that the lower the overall viral fitness, the larger the proportion of *V*_2_ among the infected population that is achieved.

**Figure 3:**
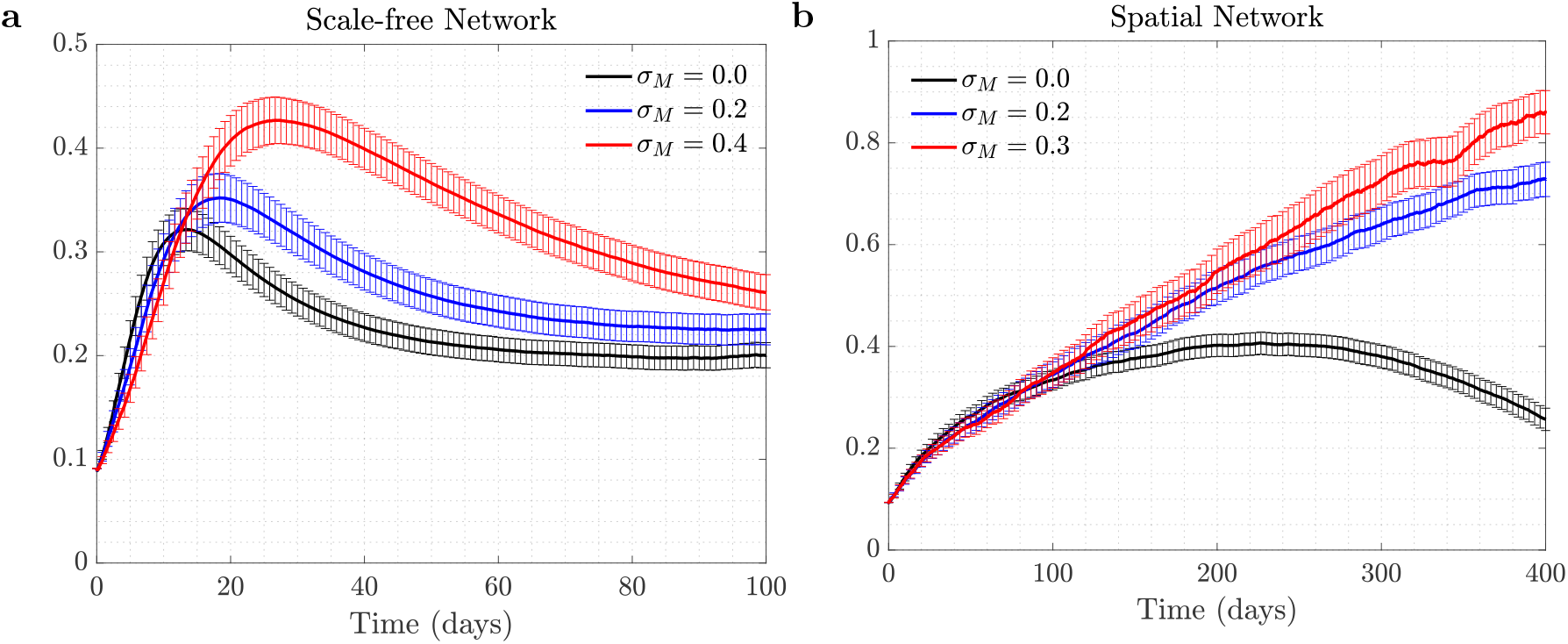
The effect of mandated SD on the proportion of *V*_2_. The proportion of the asymptomatic strain, *V*_2_, is shown as a function time, for three different levels of mandated SD: (a) Scale-free network, *σ*_*M*_ = 0 (black), *σ*_*M*_ = 0.2 (blue), and *σ*_*M*_ = 0.4, with *σ*_*S*_ = 0.4; (b) spatial network, *σ*_*M*_ = 0 (black), *σ*_*M*_ = 0.2 (blue), and *σ*_*M*_ = 0.3 (red), with *σ*_*S*_ = 0.2. All the other parameters are as in Figure 1. The levels for mandated and self-regulated SD are selected in such a way that ℛ_0_ remains above one so an outbreak for *V*_1_ is observed.

### 3.4 The effect of time-lag on *V*_2_-selection

All the simulations shown so far assumed that self-regulated SD behavior was triggered in an individual as soon as a *V*_1_-infected individual became infectious; i.e. there is no pre-symptomatic infection period and the infection status is known instantly. In reality, however, there could be a delay between a neighbor’s infection and a change in the individual’s behavior, caused by a delayed onset of symptoms, delayed testing, or a lag in information spread. Figure 4 explores the scenario where a number of days passes between an infection event and the time when self-regulated SD starts.

**Figure 4:**
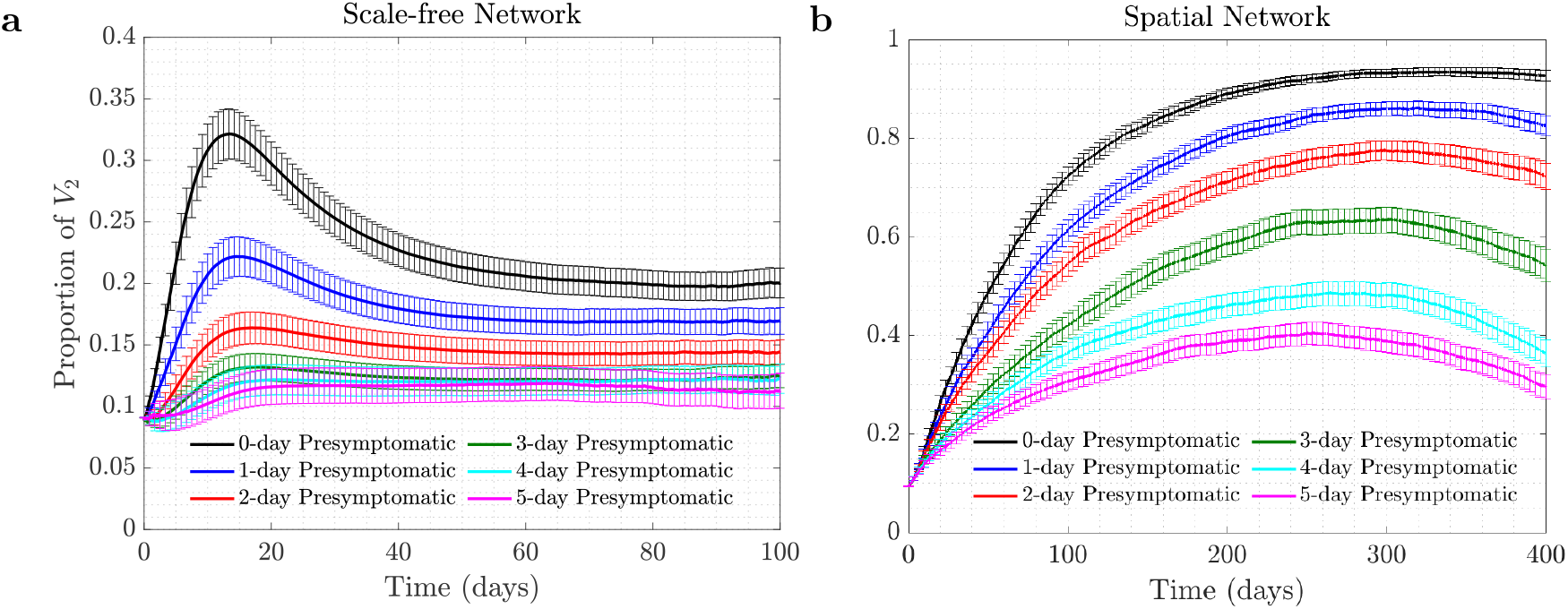
The effect of delay of self-regulated SD on selection of *V*_2_. The proportion of *V*_2_ is shown as time series for (a) scale-free and (b) spatial networks, in the presence of time delay. The different colors correspond to the time-delay of 0, 1, …, 5 days. Here *σ*_*S*_ = 0.4, *σ*_*M*_ = 0.0, and the rest of the parameters are as in figure 1.

We can see that a delay reduces positive selection experienced by the asymptomatic strain. Under scale-free networks, for the parameters in figure 4, in the presence of a 5-day lag, an increase in the fraction of *V*_2_ is almost completely eliminated. Again, because the positive selection for *V*_2_ is much stronger under spatial networks, we still observe a significant rise in the fraction of *V*_2_ in panel (b) even in the presence of a 5-day delay in protection.

### 3.5 Co-dynamics of strains on New Orleans social network

So far we have investigated the co-dynamics of viral strains on two random networks, scale-free and spatial. Both of them reflect different features of human interaction networks, but possess many very different mathematical properties. All the major results were consistent for both networks. As the next step, we will use a real-world network to demonstrate that the same trends continue to hold there.

The synthetic network that we employ here was constructed to statistically match the demographics of New Orleans residents, based on the 2009 census data. Of approximately 400, 000 residents living in 190, 000 households, the synthetic network’s sample contains 150, 000 individuals. These individuals comprise the set of network’s nodes, and the edges represent contacts of synthetic individuals through some activity types, such as “home”, “work”, “school”, “shopping”, etc. The network statistically reflects the social connections of the city’s population. Each edge of the network is labeled with one of the activity types and contains information on the amount of time spent on these contacts per day, resulting in a weighted network [53, 54]. We assumed that the amount of time of contact to cause an infection event is 15 minutes (or 0.01 of day, which is based on COVID19 infection [63]); therefore, we removed all edges with the weight less than 0.01. The resulting network has average degree 15.82 and average clustering coefficient 0.32. The degree distribution of this synthetic network is shown in figure 5. To further parameterize the model, we chose the same removal probability as in the random networks studied above, and adjusted the probability of transmission to obtain ℛ_0_ = 2.5.

**Figure 5:**
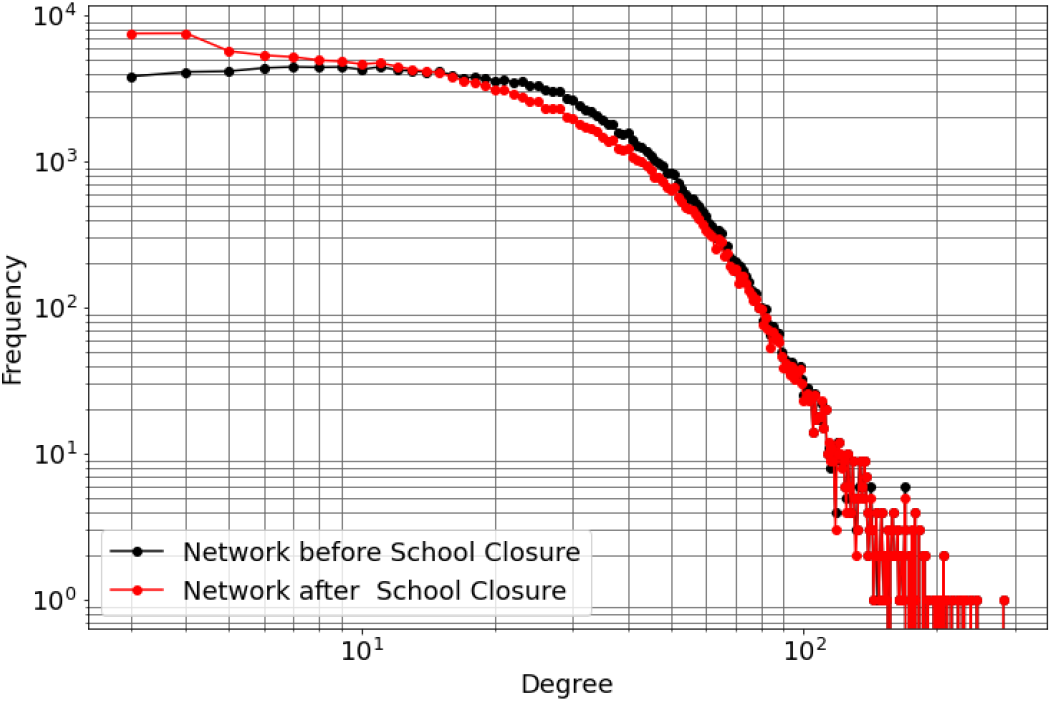
Degree distribution of the New Orleans synthetic network. Red: the basic network; black: the network under school closure (see supplementary material). The network includes 150, 000 nodes and has average degree 15.82 (with average degree 12.67 under school closure).

Figure 6 presents the time series of prevalence of the two viruses and the proportion of *V*_2_ under different levels of self-regulated SDs, in the absence of mandated SD. As established with the two types of random networks, the presence of self-regulated SD confers selective advantage to the asymptomatic virus strain, *V*_2_. We observe that self-regulated SD at level *σ*_*S*_ = 0.4 reduces the peak of the symptomatic strain, *V*_1_, to less than a half, and at level *σ*_*S*_ = 0.7 it reduces the peak of *V*_1_ by about a factor of 10, while the impact on the peak of *V*_2_ is a lot more modest. The proportion of *V*_2_ in the right panel of figure 6 increases to a peak, and this effect is stronger for higher levels of self-regulated SD. These results are consistent with those obtained for the random networks.

**Figure 6:**
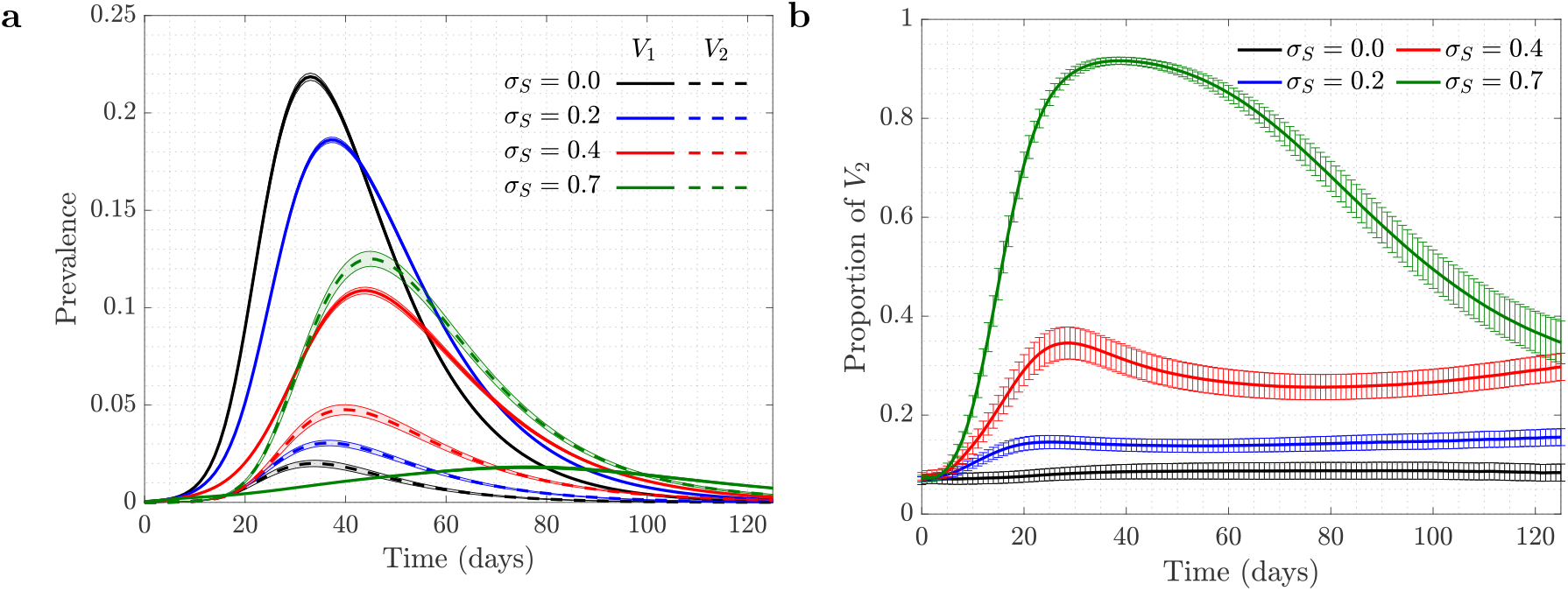
New Orleans Network of size 150, 000 individuals: the role of self-regulated SD in the spread of viruses. Time series are shown for four scenarios of no (*σ*_*S*_ = 0, black), low (*σ*_*S*_ = 0.2, blue), moderate (*σ*_*S*_ = 0.4, red), and high (*σ*_*S*_ = 0.7, green) self-regulated SD, in the absence of mandated SD. Panel (a) plot is the prevalence of *V*_1_ (solid) and *V*_2_ (dashed); panel (b) shows the proportion of *V*_2_ (*V*_2_/(*V*_1_ + *V*_2_)). *β*_1_ = *β*_2_ = 0.2 and all the other parameters are as in figure 1 (corresponding to ℛ_0_ = 2.5). Means and standard errors are shown for 1000 stochastic realizations.

Figure 7 explores the effect of mandated SD in the presence of an intermediate-level self-regulated SD, *σ*_*S*_ = 0.4. Again, the results are consistent with those observed for random networks. Increasing the level of mandated SD can make the selection for *V*_2_ significantly stronger.

**Figure 7:**
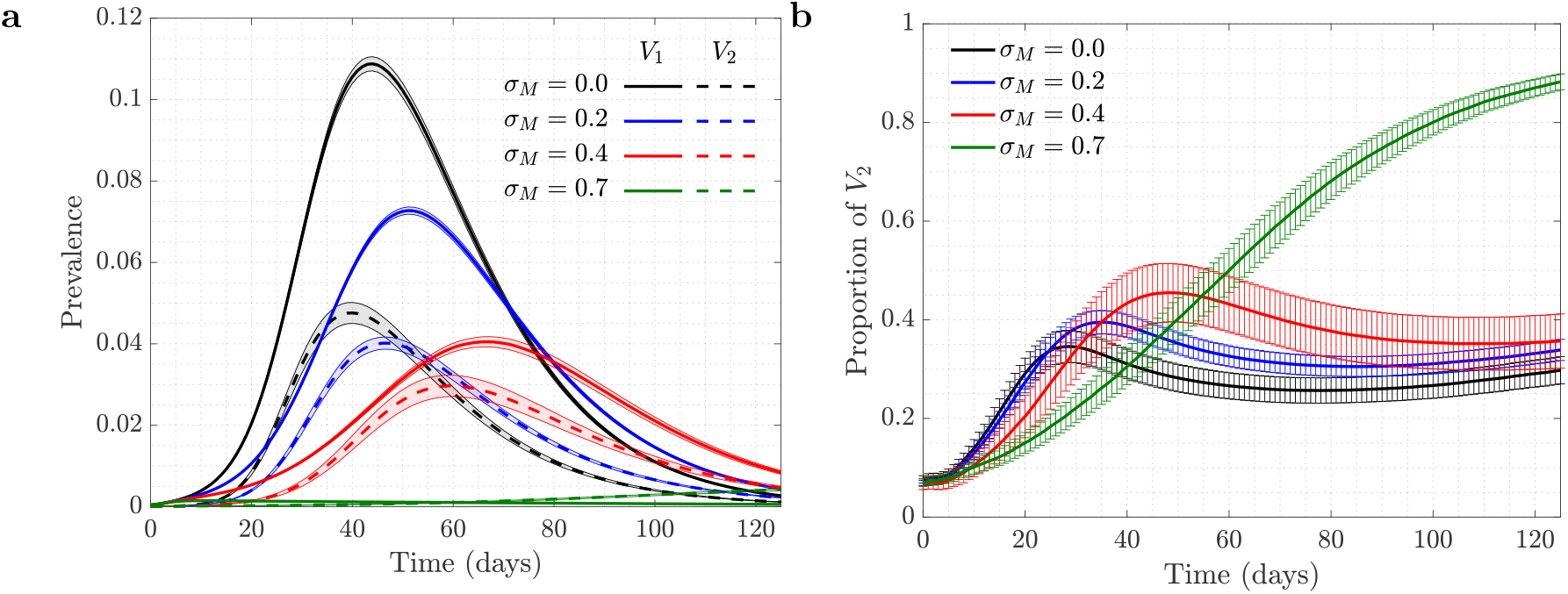
New Orleans Network of size 150, 000 individuals: the role of mandated SD in the spread of viruses. Time series are shown for four scenarios of no (*σ*_*M*_ = 0, black), low (*σ*_*M*_ = 0.2, blue), moderate (*σ*_*M*_ = 0.4, red), and high (*σ*_*M*_ = 0.6, green) mandated SD, in the presence of moderate self-regulated SD (*σ*_*S*_ = 0.4). Panel (a) plot is the prevalence of *V*_1_ (solid) and *V*_2_ (dashed); panel (b) show the proportion of *V*_2_ (*V*_2_/(*V*_1_ +*V*_2_)). *β*_1_ = *β*_2_ = 0.2 and all the other parameters are as in figure 1 (corresponding to ℛ_0_ = 2.5). Means and standard errors are shown for 1000 stochastic realizations.

## 4 Discussion

It has been reported in the literature that human behavior can change as a reaction to disease observed in others, see e.g. [64, 65, 66, 68, 69, 70]. It has further been emphasized that such behavioral changes can be an important factor in epidemic spread, e.g. in the context of sexually transmitted diseases [71, 72], or more generally [20, 21, 22, 23]. It has been noted that human behavioral traits in disease avoidance are under selection in the presence of infectious diseases [28]. Here we explore a complimentary trend: the pathogen itself might experience a force of selection to become less “visible”, or less “symptomatic”, in the presence of such human behavioral trends.

We used a discrete-time stochastic network model to investigate the spread of two co-circulating virus strains, one of which (*V*_1_) is symptomatic and the other (*V*_2_) asymptomatic. The resident strain (*V*_1_) is assumed to give rise to a mutant strain (*V*_2_) sometime during the epidemic. Three types of networks are studied: scale-free and spatial random networks, and a real-world synthetic network statistically describing social activity of individuals in New Orleans. We implemented two types of social distancing, self-regulated SD and mandated SD. Under mandated social distancing, individuals cut a given fraction of their contacts randomly, while in self-regulated social distancing, individuals opt to protect themselves based on their contacts’ infection status. More precisely, individuals cut some of their connections randomly if they find a symptomatically infected individual among their contacts.

We observed that in the presence of self-regulated protection against symptomatic cases (self-regulated SD), the proportion of asymptomatic carriers increased over time with a stronger effect corresponding to higher levels of self-regulated SD. Adding mandated SD made the effect more significant: the proportion of *V*_2_ increased for a longer duration of time and reached a higher maximum in the presence of mandated SD. Interestingly, the intensity of these trends was higher for spatial (more homogeneous and clustered) networks compared with the scale-free network, which was a result of more local infection spread and community structure. When the simulations were repeated for the real-world social network based on the New Orleans data, the selection effect was more similar to that observed for the scale-free than for the spatial network.

The selection effects observed could be weakened, e.g., by the existence of an inherent fitness disadvantage of *V*_2_ (as a result for example of a lower infectivity of this strain), or by a time-delay that exists between the onset of infection *V*_1_ and the change of behavior triggered under self-regulated SD. Nonetheless we have shown that even in the presence of these factors the selective advantage of the asymptomatic strain resulting from human behavior can still be significant and lead to a noticeable shift in the prevalence of this virus type.

While our model suggests that cautious human behavior can select for a virus variant that is less symptomatic, this selection pressure can in principle also lead to more complex outcomes. A similar advantage would be gained if the onset of symptoms was delayed and if the host could transmit the virus during this prolonged pre-symptomatic phase. Such a virus variant would also evade the behavioral reduction of network connections, yet this variant does not have to be less symptomatic or be less pathogenic. This might be at work to some extent with the SARS-CoV-2 delta variant, which is characterized by a longer window between testing positive and developing symptoms compared to previous variants [73]. Although the delta variant appears to produce higher viral loads than previous variants [74], which alone can lead to a significant transmission advantage, the longer duration of an infectious pre-symptomatic phase of delta can lead to a strong amplification of this advantage if people adjust their behavior in response to symptomatic social contacts. This might be an important contributor to the rapid rise of this variant across the globe.

The model presented here is a simplification of reality. Modeling human behavior is challenging, and here we ignored many complexities by for example assuming that individuals remove connections probabilistically when learning of a symptomatically infected individual among their circle. This approach does not distinguish between agents’ acquaintances and random contacts such as encounters in a supermarket. It also ignores demographic and socioeconomic factors that may be linked to adopting new behaviors to avoid getting infected. In addition, a static network of contacts has been assumed while in reality individuals may not have the same contacts every time unit. While further modeling efforts might address some of these shortcomings, the present model is a demonstration of principle, and not an attempt to quantitatively predict the dynamics.

Despite these uncertainties, our analysis shows robustly that human behavior in response to an infection outbreak can modulate the evolutionary trajectory of the virus. In particular, a cautious reaction of people to personal contacts that display symptomatic disease can promote the emergence of virus strains that induce less symptomatic disease. While we have not modeled one particular infection, the modeling approach is geared to describing generic respiratory infections that are transmitted through casual social contact, and therefore has implications for the current SARS-CoV-2 pandemic.

## Appendix

### ODE modeling

SIR models based on ordinary differential equations are an important tool in epidemiological infection studies [67], and they have been widely used for various emerging infections such as COVID19 [75]. Here we denote by *x* the fraction of susceptible individuals, and distinguish between two strains of infection, *V*_1_ and *V*_2_. The fraction of individuals infected with *V*_1_ is denoted by *y*_1_ and the fraction of individuals infected with *V*_2_ is denoted by *y*_2_. We assume that an individual cannot be super-infected with a different virus, and that recovered individuals cannot be infected anymore. This gives rise to the following system:

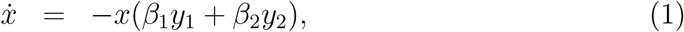

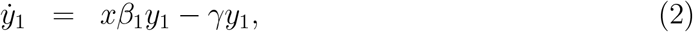

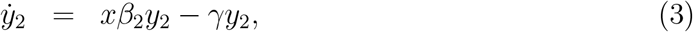

with initial conditions

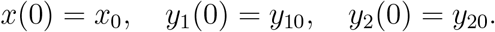

Here *β*_1_ and *β*_2_ are the rate of infection for the two strains, and *γ* the rate of removal. Let us denote by *z* the proportion of the individals infected with *V*_2_:

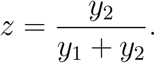

This quantity satisfies the following equation:

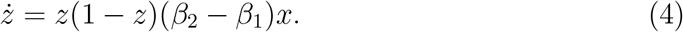

In particular, if the two strains are neutral to each other (*β*_1_ = *β*_2_) then the fraction *z* is expected to stay constant. It will increase if *β*_2_ > *β*_1_ and decrease if *β*_2_ < *β*_1_. Let us consider the problem where *V*_2_ is an advantageous mutant (*β*_2_ > *β*_1_), which is initially in a minority, that is, *y*_20_ ≪ *y*_10_. We note that in this case, *z* will be an increasing function of time. Its initial growth is exponential with the rate approximately given by *β*_2_ – *β*_1_ (assuming that *x* ≈ *x*_0_ ≈ 1). As *x* decreases, the growth slows down. Two extreme scenarios can be distinguished, see figure 8:

**Figure 8:**
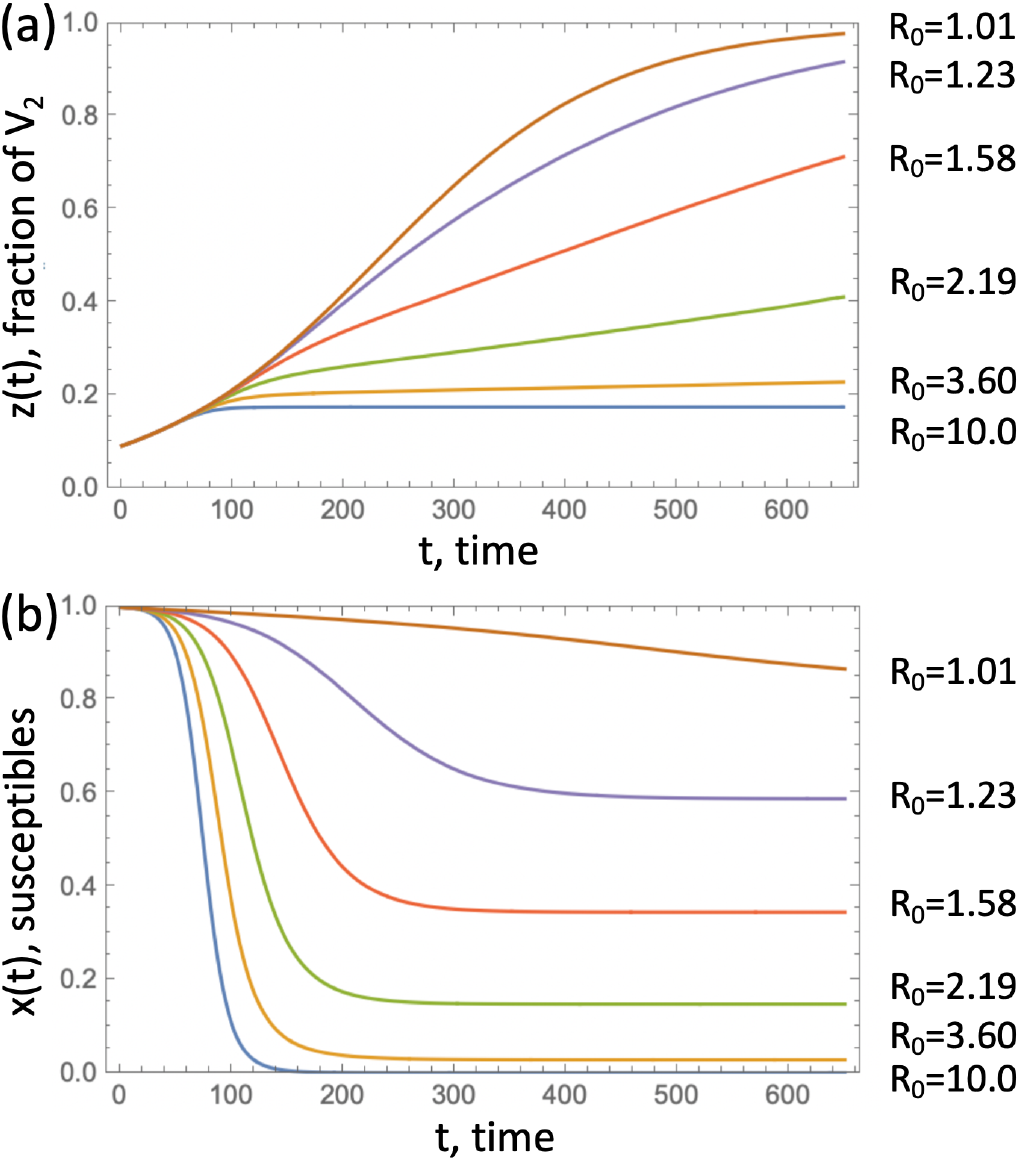
The fraction of an advantageous virus, *V*_2_. (a) The quantity *z*(*t*) obtained by solving equations (1-3) is plotted as a function of time for several values of *R*_0_, obtained by changing the death rate, *a*. (b) The corresponding susceptible populations as functions of time. The rest of the parameters are *β*_1_ = 0.1, *β*_2_ = 0.11, *y*_1_(0) = 0.001, *y*_2_(0) = 0.1*y*_1_(0).

1. *z* approaches 1 well before *x* decreases significantly; in this case the dynamics of *z* is well described by the logistic growth model.
2. The epidemic ends well before *z* approaches 1, in which case near the epidemic end, the growth of *z* becomes linear with the rate approximately given by (*β*_2_ − *β*_1_)*x*_∞_, where 1 − *x*_∞_ is the final epidemic size.

We observe that larger overall values of *R*_0_ correspond to a more modest expansion of the advantageous virus *V*_2_ (assuming that the % advantage is fixed; it is for example 10% in figure 8).

In this context, it is useful to calculate the value

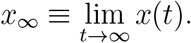

If *β*_2_ = *β*_1_, the we have the following final size relation:

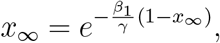

which is an implicit formula for *x*_∞_. In the case of two different pathogens, if we denote 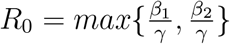, we have [76]

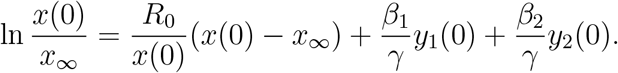

The ODE model can be used to calculate the proportion of *V*_2_ by the end of the epidemic. Figure 9 shows an example where we fixed the values *β*_1_ and *β*_2_, such that *V*_2_ has a 10% advantage in terms of infectivity, and also assumed that *y*_2_(0) = 0.1*y*_1_(0). Parameters *γ* and *y*_1_(0) were varied over a wide range, which corresponds to varying *R*_0_ (associated with the resident virus) and the total population size relative to the initial number of infected individuals. Panel (a) illustrates the way we numerically calculate the end of epidemic time, *t*_end_, and panel (b) shows the fraction of *V*_2_ at time *t*_end_ as a function of *R*_0_ and log_10_ *y*_1_(0).

**Figure 9:**
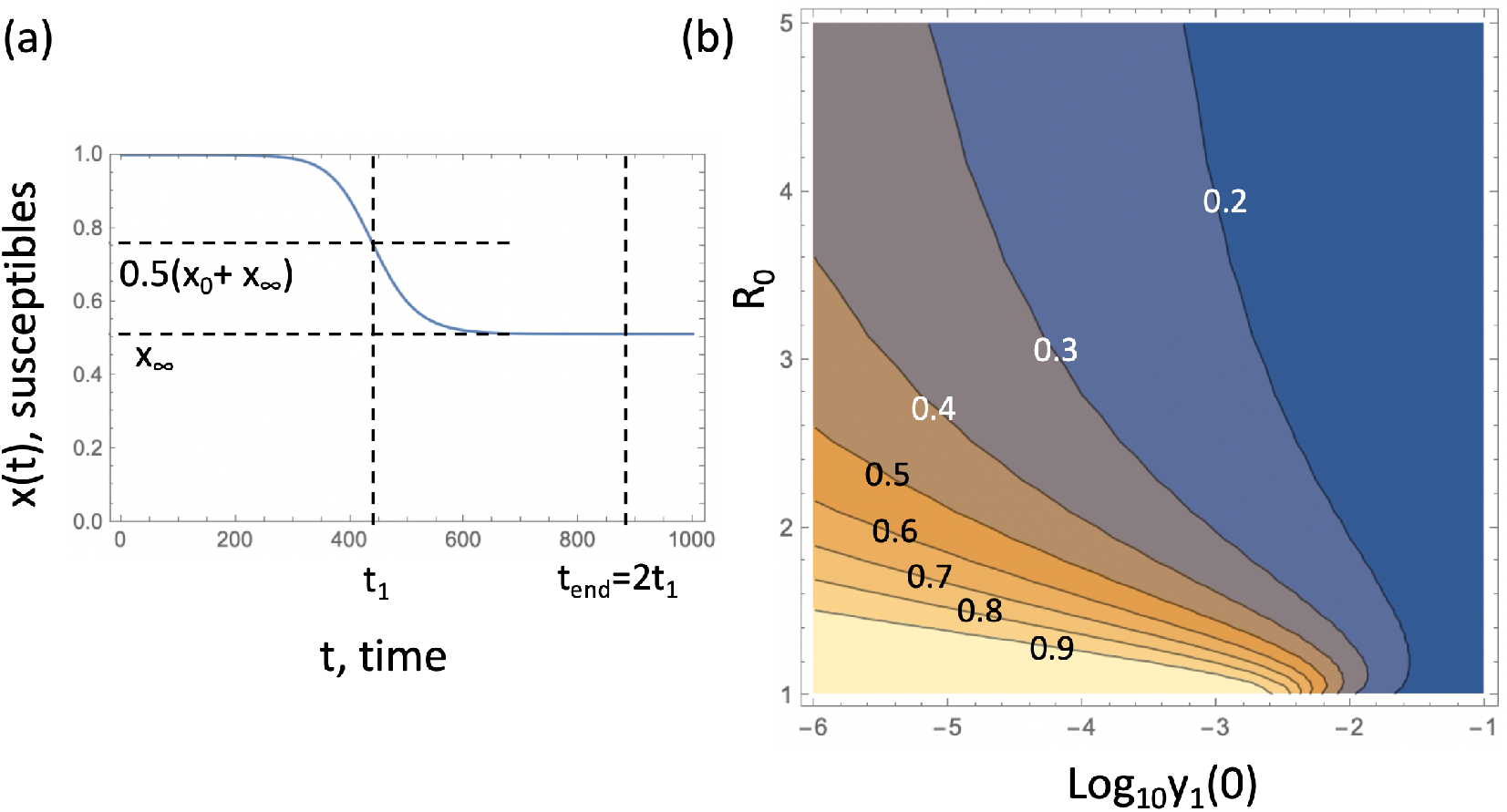
Fraction of *V*_2_ at the end of the epidemic. (a) Calculation of *t*_end_, which represents the end of the epidemic, is illustrated. The blue line is the fraction of susceptible individuals, *x*(*t*), obtained as a solution of equations (1-3); *t*_end_ = 2*t*_1_, where *t*_1_ corresponds to 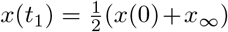. In other words, at time *t*_1_ the population of susceptible individuals reaches halfway to its final value, *x*_∞_. (b) Quantity *y*_2_/(*y*_1_ + *y*_2_) obtained by solving equations (1-3), is plotted at time *t*_end_, as a function of the initial proportion of individuals infected with *V*_1_, and *R*_0_. The rest of the parameters are *β*_1_ = 0.1, *β*_2_ = 0.11, *y*_2_(0) = 0.1*y*_1_(0).

We observe that typically, increasing *R*_0_ leads to a smaller final fraction of *V*_2_. For relatively large *R*_0_ values, the fraction of susceptible individuals decreases quickly leading to an extremely slow linear growth of the fraction *z*(*t*). On the other hand, decreasing *y*_1_(0) (which is equivalent to considering larger total populations) leads to an increase in the final fraction of *V*_2_. Larger populations result in a longer epidemic, and *V*_2_ consequently has a longer time to gain on *V*_1_.

### Further details of viral co-dynamics

In figure 1 in the main text, as well as others (such as figures 2 and 3 in the main text), we observe that the fraction of *V*_2_ often has a one-humped shape: it first increases to a peak and then decreases as the epidemic dwindles down. This is a phenomenon that does not have an analogy in the simple ODE model, (1-3). Equation (4) for the fraction suggests that the proportion of *V*_2_ always increases if *β*_2_ > *β*_1_. On the other hand, in the agent-based models for symptomatic virus *V*_1_ and its asymptomatic counterpart, *V*_2_, we observe that, both for scale-free and spatial networks, the numerical gain of *V*_2_ eventually decreases. This is related to the epidemic duration of the two strands: the advantageous virus experiences a shorter epidemic, and this effect increases with the amount of advantage. Figure 10 shows that the time it take *V*_2_ to reach its infection peak is shorter compared to that for *V*_1_, and as we increase the level of self-regulated SD (thus increasing the advantage of *V*_2_), the difference in the peak time grows. Therefore, there is a time-interval during which the amount of *V*_2_ infection already decreases while *V*_1_ still grows towards its peak, resulting in a reduction in the *V*_2_ fraction.

**Figure 10:**
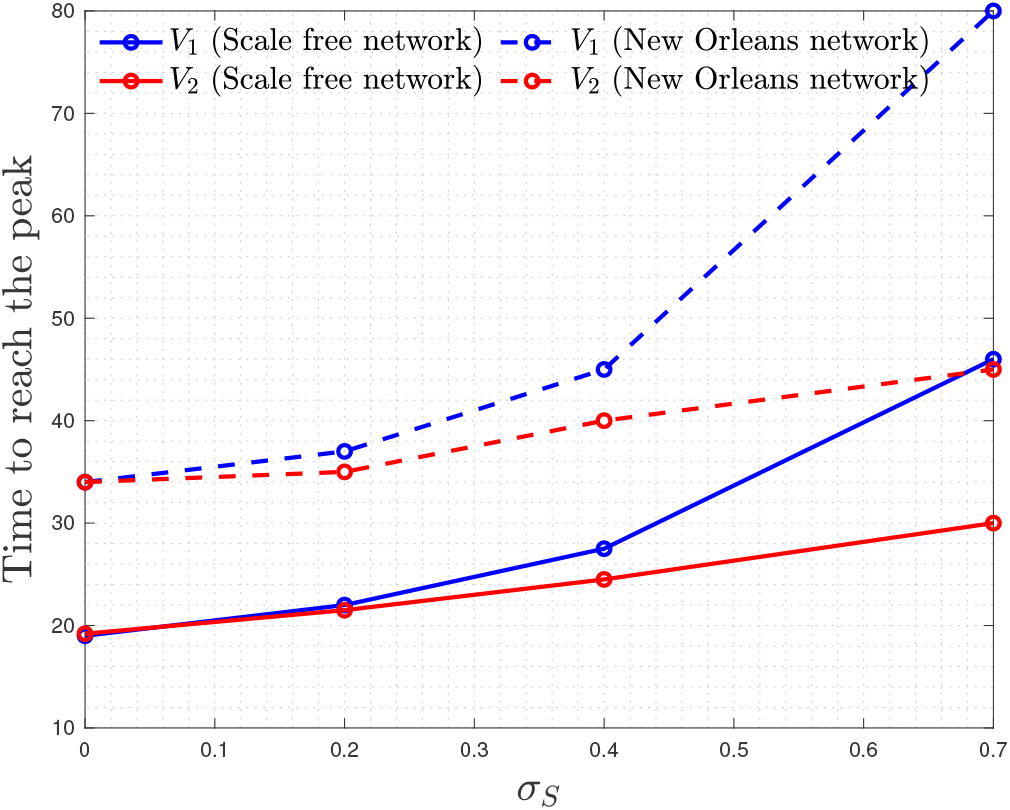
Time to reach infection peak for viruses *V*_1_ (blue) and *V*_2_(red), as a function of *σ*_*S*_ (the measure of *V*_2_ advantage).

Note that this is not observed in the ODE system and also was less pronounced in more clustered spatial network. In ODE model, the peak of infection *y*_*i*_ is reached when 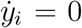, which corresponds to the time *t*_*i*_ when 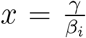, for *i* ∈ {1, 2}. Since *x*(*t*) is a decreasing function and *β*_2_ > *β*_1_ (in analogy with self-regulated SD), we necessarily conclude that *t*_2_ > *t*_1_, that is, the epidemic corresponding to a more infectious type is always longer.

### Co-dynamics of strains on New Orleans social network under school closure

School closure is an important component of social distancing measures, which has for example been implemented widely during the SARS-CoV-2 pandemic. Therefore, we have repeated the analysis of Section 3.5 in the main text after removing all the edges related to “school”. Degree distribution of the resulting network is shown in black in figure 5 in the main text. We tuned up the transmission rates *β*_1_ and *β*_2_ to have ℛ_0_ = 2.5 and reran our simulations on the new network, see figures 11 and 12.

**Figure 11:**
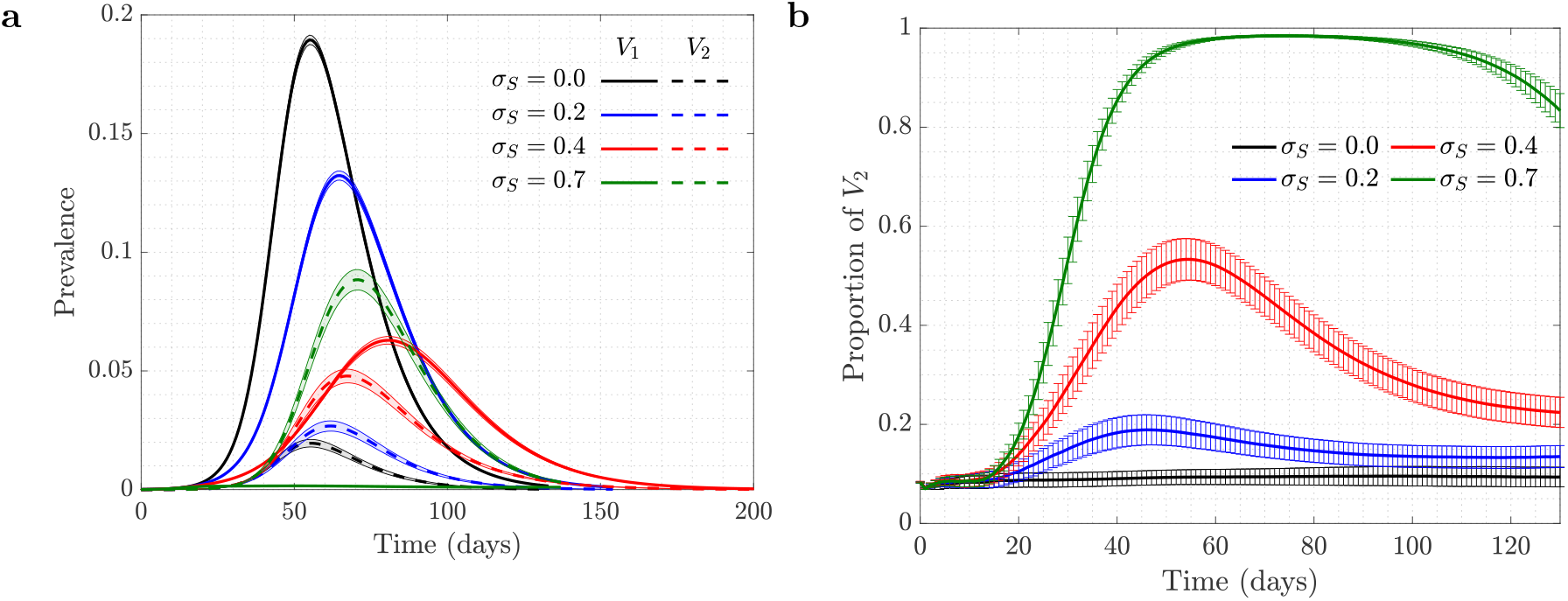
New Orleans Network under school closure: the role of self-regulated SD in the spread of viruses. Time series are shown for four scenarios of no (*σ*_*S*_ = 0, black), low (*σ*_*S*_ = 0.2, blue), moderate (*σ*_*S*_ = 0.4, red), and high (*σ*_*S*_ = 0.7, green) self-regulated SD, in the absence of mandated SD. Panel (a) plot is the prevalence of *V*_1_ (solid) and *V*_2_ (dashed); panel (b) shows the proportion of *V*_2_ (*V*_2_/(*V*_1_ + *V*_2_)). *β*_1_ = *β*_2_ = 0.29 and all the other parameters are as in figure 1 in the main text (corresponding to ℛ_0_ = 2.5). Means and standard errors are shown for 1000 stochastic realizations.

**Figure 12:**
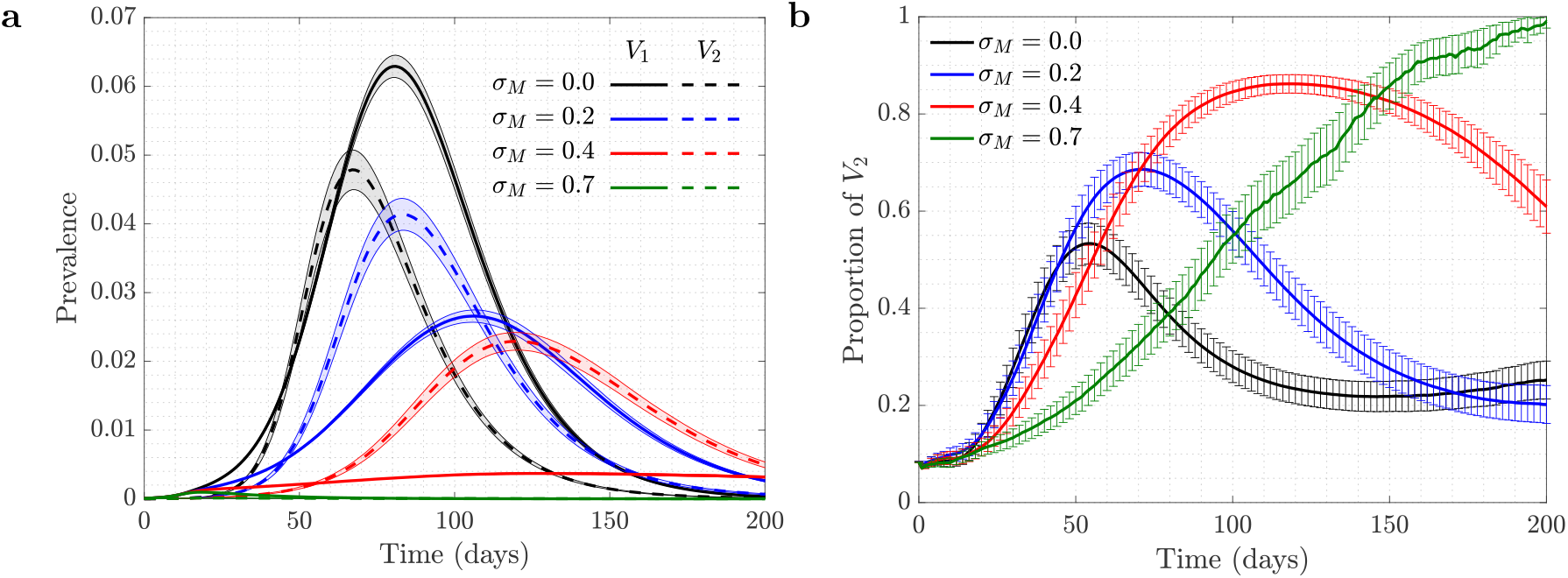
New Orleans Network under school closure: the role of mandated SD in the spread of viruses. Time series are shown for four scenarios of no (*σ*_*M*_ = 0, black), low (*σ*_*M*_ = 0.2, blue), moderate (*σ*_*M*_ = 0.4, red), and high (*σ*_*M*_ = 0.6, green) mandated SD, in the presence of moderate self-regulated SD (*σ*_*S*_ = 0.4). Panel (a) plot is the prevalence of *V*_1_ (solid) and *V*_2_ (dashed); panel (b) show the proportion of *V*_2_ (*V*_2_/(*V*_1_ + *V*_2_)). *β*_1_ = *β*_2_ = 0.29 and all the other parameters are as in figure 1 in the main text (corresponding to ℛ_0_ = 2.5). Means and standard errors are shown for 1000 stochastic realizations.

In figure 11, we implemented the impact of various levels for self-regulated SD (*σ*_*S*_ = 0, 0.2, 0.4, and 0.7) in the absence of mandated SD (*σ*_*M*_ = 0). Similar to previous results, increasing the level of self-regulated SD causes more selective advantage to the asymptomatic virus strain, *V*_2_.

Figure 12 explores the effect of mandated SD in the presence of an intermediate-level self-regulated SD, *σ*_*S*_ = 0.4. Again, and similar to the results of Section 3.5 in the main text, increasing the level of mandated SD causes that the selection for *V*_2_ to become significantly stronger.

While the results for the New Orleans Network are qualitatively similar with and without school closure, we notice that the effect of further SD measures on the background of closed schools is stronger, since we start with a somewhat sparser network.

## References

[1] Stefan Ma and Yingcun Xia. Mathematical understanding of infectious disease dynamics, volume 16. World Scientific, 2009.

[2] Odo Diekmann, Hans Heesterbeek, and Tom Britton. Mathematical tools for understanding infectious disease dynamics, volume 7. Princeton University Press, 2012.

[3] Constantinos I Siettos and Lucia Russo. Mathematical modeling of infectious disease dynamics. Virulence, 4(4):295–306, 2013.

[4] Hans Heesterbeek, Roy M Anderson, Viggo Andreasen, Shweta Bansal, Daniela De Angelis, Chris Dye, Ken TD Eames, W John Edmunds, Simon DW Frost, Sebastian Funk, et al. Modeling infectious disease dynamics in the complex landscape of global health. Science, 347(6227), 2015.

[5] Sarah Cobey. Modeling infectious disease dynamics. Science, 368(6492):713– 714, 2020.

[6] Zhilan Feng, Ulf Dieckmann, and Simon A Levin. Disease evolution: models, concepts, and data analyses, volume 71. American Mathematical Soc., 2006.

[7] Troy Day, Samuel Alizon, and Nicole Mideo. Bridging scales in the evolution of infectious disease life histories: theory. Evolution: International Journal of Organic Evolution, 65(12):3448–3461, 2011.

[8] Chiara Poletto, Sandro Meloni, Vittoria Colizza, Yamir Moreno, and Alessandro Vespignani. Host mobility drives pathogen competition in spatially structured populations. PLoS Comput Biol, 9(8):e1003169, 2013.

[9] Quentin Griette, Gaël Raoul, and Sylvain Gandon. Virulence evolution at the front line of spreading epidemics. Evolution, 69(11):2810–2819, 2015.

[10] Chiara Poletto, Sandro Meloni, Ashleigh Van Metre, Vittoria Colizza, Yamir Moreno, and Alessandro Vespignani. Characterising two-pathogen competition in spatially structured environments. Scientific reports, 5(1):1–9, 2015.

[11] Sylvain Gandon, Troy Day,C Jessica E Metcalf, and Bryan T Grenfell. Forecasting epidemiological and evolutionary dynamics of infectious diseases. Trends in ecology & evolution, 31(10):776–788, 2016.

[12] Erik E Osnas, Paul J Hurtado, and Andrew P Dobson. Evolution of pathogen virulence across space during an epidemic. The American Naturalist, 185(3):332–342, 2015.

[13] Mark EJ Newman. Threshold effects for two pathogens spreading on a network. Physical review letters, 95(10):108701, 2005.

[14] Brian Karrer and Mark EJ Newman. Competing epidemics on complex networks. Physical Review E, 84(3):036106, 2011.

[15] Shweta Bansal and Lauren Ancel Meyers. The impact of past epidemics on future disease dynamics. Journal of theoretical biology, 309:176–184, 2012.

[16] Joel C Miller. Cocirculation of infectious diseases on networks. Physical Review E, 87(6):060801, 2013.

[17] Gabriel E Leventhal, Alison L Hill, Martin A Nowak, and Sebastian Bonhoeffer. Evolution and emergence of infectious diseases in theoretical and real-world networks. Nature communications, 6(1):1–11, 2015.

[18] Sébastien Lion and Sylvain Gandon. Spatial evolutionary epidemiology of spreading epidemics. Proceedings of the Royal Society B: Biological Sciences, 283(1841):20161170, 2016.

[19] Francesco Pinotti, Éric Fleury, Didier Guillemot, Pierre-Yves Böelle, and Chiara Poletto. Host contact dynamics shapes richness and dominance of pathogen strains. PLoS computational biology, 15(5):e1006530, 2019.

[20] Sebastian Funk, Marcel Salathé, and Vincent AA Jansen. Modelling the influence of human behaviour on the spread of infectious diseases: a review. Journal of the Royal Society Interface, 7(50):1247–1256, 2010.

[21] Ceyhun Eksin, Jeff S Shamma, and Joshua S Weitz. Disease dynamics in a stochastic network game: a little empathy goes a long way in averting outbreaks. Scientific reports, 7(1):1–13, 2017.

[22] Ceyhun Eksin, Keith Paarporn, and Joshua S Weitz. Systematic biases in disease forecasting–the role of behavior change. Epidemics, 27:96–105, 2019.

[23] Joshua S Weitz, Sang Woo Park, Ceyhun Eksin, and Jonathan Dushoff. Awareness-driven behavior changes can shift the shape of epidemics away from peaks and toward plateaus, shoulders, and oscillations. Proceedings of the National Academy of Sciences, 117(51):32764–32771, 2020.

[24] Samuel V Scarpino, Antoine Allard, and Laurent Hébert-Dufresne. The effect of a prudent adaptive behaviour on disease transmission. Nature Physics, 12(11):1042–1046, 2016.

[25] Lorna Fewtrell, Rachel B Kaufmann, David Kay, Wayne Enanoria, Laurence Haller, and John M Colford Jr. Water, sanitation, and hygiene interventions to reduce diarrhoea in less developed countries: a systematic review and meta-analysis. The Lancet infectious diseases, 5(1):42–52, 2005.

[26] Chris T Bauch and David JD Earn. Vaccination and the theory of games. Proceedings of the National Academy of Sciences, 101(36):13391–13394, 2004.

[27] Sara Del Valle, Herbert Hethcote, James M Hyman, and Carlos Castillo-Chavez. Effects of behavioral changes in a smallpox attack model. Mathematical biosciences, 195(2):228–251, 2005.

[28] Mark M Tanaka, Jochen Kumm, and Marcus W Feldman. Coevolution of pathogens and cultural practices: a new look at behavioral heterogeneity in epidemics. Theoretical population biology, 62(2):111–119, 2002.

[29] Joshua M Epstein, Jon Parker, Derek Cummings, and Ross A Hammond. Coupled contagion dynamics of fear and disease: mathematical and computational explorations. PLoS One, 3(12):e3955, 2008.

[30] Thilo Gross, Carlos J Dommar D’Lima, and Bernd Blasius. Epidemic dynamics on an adaptive network. Physical review letters, 96(20):208701, 2006.

[31] Sebastian Funk, Erez Gilad, Chris Watkins, and Vincent AA Jansen. The spread of awareness and its impact on epidemic outbreaks. Proceedings of the National Academy of Sciences, 106(16):6872–6877, 2009.

[32] KM Ariful Kabir, Kazuki Kuga, and Jun Tanimoto. Effect of information spreading to suppress the disease contagion on the epidemic vaccination game. Chaos, Solitons & Fractals, 119:180–187, 2019.

[33] Jia-Qian Kan and Hai-Feng Zhang. Effects of awareness diffusion and self-initiated awareness behavior on epidemic spreading-an approach based on multiplex networks. Communications in Nonlinear Science and Numerical Simulation, 44:193–203, 2017.

[34] Chengjun Sun, Wei Yang, Julien Arino, and Kamran Khan. Effect of media-induced social distancing on disease transmission in a two patch setting. Mathematical biosciences, 230(2):87–95, 2011.

[35] Zhen Wang, Michael A Andrews, Zhi-Xi Wu, Lin Wang, and Chris T Bauch. Coupled disease–behavior dynamics on complex networks: A review. Physics of life reviews, 15:1–29, 2015.

[36] Qingchu Wu, Xinchu Fu, Michael Small, and Xin-Jian Xu. The impact of awareness on epidemic spreading in networks. Chaos: an interdisciplinary journal of nonlinear science, 22(1):013101, 2012.

[37] Robert J Glass, Laura M Glass, Walter E Beyeler, and H Jason Min. Targeted social distancing designs for pandemic influenza. Emerging infectious diseases, 12(11):1671, 2006.

[38] LD Valdez, Pablo A Macri, and Lidia A Braunstein. Intermittent social distancing strategy for epidemic control. Physical Review E, 85(3):036108, 2012.

[39] Joseph A Lewnard and Nathan C Lo. Scientific and ethical basis for socialdistancing interventions against covid-19. The Lancet. Infectious diseases, 20(6):631, 2020.

[40] Kai Kupferschmidt and Jon Cohen. Can China’s COVID-19 strategy work elsewhere? Science, 367(6482):1061–1062, 2020.

[41] Zhilan Feng, John W Glasser, and Andrew N Hill. On the benefits of flattening the curve: A perspective. Mathematical Biosciences, 326:108389, 2020.

[42] Zoltán Vokó and János György Pitter. The effect of social distance measures on covid-19 epidemics in europe: an interrupted time series analysis. GeroScience, 42(4):1075–1082, 2020.

[43] Rajiv Chowdhury, Shammi Luhar, Nusrat Khan, Sohel Reza Choudhury, Imran Matin, and Oscar H Franco. Long-term strategies to control covid-19 in low and middle-income countries: an options overview of community-based, nonpharmacological interventions. European journal of epidemiology, 35(8):743– 748, 2020.

[44] Stephen M Kissler, Christine Tedijanto, Marc Lipsitch, and Yonatan Grad. Social distancing strategies for curbing the covid-19 epidemic. medRxiv, 2020.

[45] Neil Ferguson, Daniel Laydon, Gemma Nedjati Gilani, Natsuko Imai, Kylie Ainslie, Marc Baguelin, Sangeeta Bhatia, Adhiratha Boonyasiri, ZULMA Cucunuba Perez, Gina Cuomo-Dannenburg, et al. Report 9: Impact of nonpharmaceutical interventions (npis) to reduce covid19 mortality and healthcare demand. 2020.

[46] Joel R Koo, Alex R Cook, Minah Park, Yinxiaohe Sun, Haoyang Sun, Jue Tao Lim, Clarence Tam, and Borame L Dickens. Interventions to mitigate early spread of sars-cov-2 in singapore: a modelling study. The Lancet Infectious Diseases, 2020.

[47] Joel Hellewell, Sam Abbott, Amy Gimma, Nikos I Bosse, Christopher I Jarvis, Timothy W Russell, James D Munday, Adam J Kucharski, W John Edmunds, Fiona Sun, et al. Feasibility of controlling covid-19 outbreaks by isolation of cases and contacts. The Lancet Global Health, 2020.

[48] Marino Gatto, Enrico Bertuzzo, Lorenzo Mari, Stefano Miccoli, Luca Carraro, Renato Casagrandi, and Andrea Rinaldo. Spread and dynamics of the covid-19 epidemic in italy: Effects of emergency containment measures. Proceedings of the National Academy of Sciences, 117(19):10484–10491, 2020.

[49] Chaolong Wang, Li Liu, Xingjie Hao, Huan Guo, Qi Wang, Jiao Huang, Na He, Hongjie Yu, Xihong Lin, An Pan, et al. Evolving epidemiology and impact of non-pharmaceutical interventions on the outbreak of coronavirus disease 2019 in wuhan, china. MedRxiv, 2020.

[50] Congying Liu, Xiaoqun Wu, Riuwu Niu, Xiuqi Wu, and Ruguo Fan. A new sair model on complex networks for analysing the 2019 novel coronavirus (covid-19). Nonlinear Dynamics, pages 1–11, 2020.

[51] Tom Li, Yan Liu, Man Li, Xiaoning Qian, and Susie Y Dai. Mask or no mask for covid-19: A public health and market study. PloS one, 15(8):e0237691, 2020.

[52] He Huang, Yahong Chen, and Zhijun Yan. Impacts of social distancing on the spread of infectious diseases with asymptomatic infection: A mathematical model. Applied Mathematics and Computation, 398:125983, 2021.

[53] Stephen Eubank, Christopher Barrett, Richard Beckman, Keith Bisset, Lisa Durbeck, Christopher Kuhlman, Bryan Lewis, Achla Marathe, Madhav Marathe, and Paula Stretz. Detail in network models of epidemiology: are we there yet? Journal of biological dynamics, 4(5):446–455, 2010.

[54] S Eubank. Synthetic data products for societal infrastructures and protopopulations: Data set 2.0. Technical report, Technical Report NDSSL-TR-07-003, Network Dynamics and Simulation Science Laboratory, Virginia Polytechnic Institute and State University, 2008.

[55] Jesper Dall and Michael Christensen. Random geometric graphs. Physical review E, 66(1):016121, 2002.

[56] John C Lang, Hans De Sterck, Jamieson L Kaiser, and Joel C Miller. Analytic models for SIR disease spread on random spatial networks. Journal of Complex Networks, 6(6):948–970, 2018.

[57] Albert-Laszló Barabási and Eric Bonabeau. Scale-free networks. Scientific american, 288(5):60–69, 2003.

[58] Albert-Laszló Barabási and Réka Albert. Emergence of scaling in random networks. science, 286(5439):509–512, 1999.

[59] Isiaah Crawford. Attitudes of undergraduate college students toward AIDS. Psychological Reports, 66(1):11–16, 1990.

[60] Gerardo Chowell, James M Hyman, Stephen Eubank, and Carlos Castillo-Chavez. Scaling laws for the movement of people between locations in a large city. Physical Review E, 68(6):066102, 2003.

[61] Aric Hagberg, Dan Schult, Pieter Swart, D Conway, L Séguin-Charbonneau, C Ellison, B Edwards, and J Torrents. Networkx. High productivity software for complex networks. Webová strá nka https://networkx.lanl.gov/wiki, 2004.

[62] Benjamin Roche, John M Drake, and Pejman Rohani. An agent-based model to study the epidemiological and evolutionary dynamics of influenza viruses. BMC bioinformatics, 12(1):87, 2011.

[63] Matt J Keeling, T Deirdre Hollingsworth, and Jonathan M Read. Efficacy of contact tracing for the containment of the 2019 novel coronavirus (covid-19). J Epidemiol Community Health, 74(10):861–866, 2020.

[64] A Andersson-Ellström, Lars Forssman, and Ian Milsom. The relationship between knowledge about sexually transmitted diseases and actual sexual behaviour in a group of teenage girls. Sexually Transmitted Infections, 72(1):32– 36, 1996.

[65] William W Darrow. Health education and promotion for std prevention: lessons for the next millennium. Sexually Transmitted Infections, 73(2):88–94, 1997.

[66] Jo N Hays. Epidemics and pandemics: their impacts on human history. Abcclio, 2005.

[67] Anderson, Roy M and May Robert M. Infectious diseases of humans: dynamics and control. 1992.

[68] Valerie A Curtis. Infection-avoidance behaviour in humans and other animals. Trends in Immunology, 35(10):457–464, 2014.

[69] Asma Azizi, Cesar Montalvo, Baltazar Espinoza, Yun Kang, and Carlos Castillo-Chavez. Epidemics on networks: Reducing disease transmission using health emergency declarations and peer communication. Infectious Disease Modelling, 5:12–22, 2020.

[70] Natalia L Komarova, Asma Azizi, and Dominik Wodarz. Network models and the interpretation of prolonged infection plateaus in the covid19 pandemic. Epidemics, 35:100463, 2021.

[71] Karl P Hadeler and Carlos Castillo-Chávez. A core group model for disease transmission. Mathematical biosciences, 128(1-2):41–55, 1995.

[72] James M Hyman and Jia Li. Behavior changes in sis std models with selective mixing. SIAM Journal on Applied Mathematics, 57(4):1082–1094, 1997.

[73] Min Kang, Hualei Xin, Jun Yuan, Sheikh Taslim xAli, Zimian Liang, Jiayi Zhang, Ting Hu, Eric Lau, Yingtao Zhang, Meng Zhang, et al. Transmission dynamics and epidemiological characteristics of delta variant infections in china. medRxiv, 2021.

[74] Baisheng Li, Aiping Deng, Kuibiao Li, Yao Hu, Zhencui Li, Qianling Xiong, Zhe Liu, Qianfang Guo, Lirong Zou, Huan Zhang, et al. Viral infection and transmission in a large well-traced outbreak caused by the delta sars-cov-2 variant. MedRxiv, 2021.

[75] Fang, Yaqing and Nie Yiting and Penny, Marshare Transmission dynamics of the COVID-19 outbreak and effectiveness of government interventions: A data-driven analysis. Journal of medical virology, 92(6):645–659, 2020.

[76] Arino, Julien and Brauer, Fred and van den Driessche, Pauline and Watmough, James and Wu, Jianhong. A final size relation for epidemic models. Mathematical Biosciences & Engineering, 4(2), 2007.

